# Fast estimation of L1-regularized linear models in the mass-univariate setting

**DOI:** 10.1101/2020.01.16.909234

**Authors:** Holger Mohr, Hannes Ruge

## Abstract

In certain modeling approaches, activation analyses of task-based fMRI data can involve a relatively large number of predictors. For example, in the encoding model approach, complex stimuli are represented in a high-dimensional feature space, resulting in design matrices with many predictors. Similarly, single-trial models and finite impulse response models may also encompass a large number of predictors. In settings where only few of those predictors are expected to be informative, a sparse model fit can be obtained via L1-regularization. However, estimating L1-regularized models requires an iterative fitting procedure, which considerably increases computation time compared to estimating unregularized or L2-regularized models, and complicates the application of L1-regularization on whole-brain data and large sample sizes. Here we provide several functions for estimating L1-regularized models that are optimized for the mass-univariate analysis approach. The package includes a parallel implementation of the coordinate descent algorithm for CPU-only systems and two implementations of the alternating direction method of multipliers algorithm requiring a GPU device. While the core algorithms are implemented in C++/CUDA, data input/output and parameter settings can be conveniently handled via Matlab. The CPU-based implementation is highly memory-efficient and provides considerable speed-up compared to the standard implementation not optimized for the mass-univariate approach. Further acceleration can be achieved on systems equipped with a CUDA-enabled GPU. Using the fastest GPU-based implementation, computation time for whole-brain estimates can be reduced from 9 hours to 5 minutes in an exemplary data setting. Overall, the provided package facilitates the use of L1-regularization for fMRI activation analyses and enables an efficient employment of L1-regularization on whole-brain data and large sample sizes.

## Introduction

Over the last two decades, various fMRI activation analysis approaches have been established that involve a relatively large number of predictors. For example, single-trial models can be employed to obtain activation estimates for individual experimental trials (Mumford et al., 2012). The single-trial estimates can be subsequently used as input for further analyses such as multivariate pattern analyses or connectivity analyses (Mumford et al., 2014; Rissman et al., 2004). As single-trial models require a separate predictor for each experimental trial, the resulting design matrices typically encompass a relatively large number of predictors.

The finite impulse response (FIR) model is another example for an fMRI activation analysis approach involving many predictors (Ollinger et al., 2001). In the FIR approach, the fMRI signal is modeled by a set of impulse response predictors, instead of a predefined hemodynamic response shape. This modeling approach can capture activation dynamics deviating from the canonical response shape, but typically involves a large number of predictors (growing proportionally with the size of the FIR basis set). Both in single-trial and FIR models, the large number of predictors can reduce the robustness of the beta estimates.

Another fMRI modeling approach typically involving a large number of predictors is the so-called encoding model approach (van Gerven, 2017). In this approach, stimuli are represented in a high-dimensional feature space instead of being assigned to a low-dimensional set of categories. For example, instead of assigning visual stimuli to object categories such as houses, trees, etc., the stimuli are represented by a high-dimensional vector of feature weights. Such features may comprise a set of Gabor wavelets (Kay et al., 2008), may be extracted from deep neural networks (Güclü and van Gerven, 2015) or may simply consist of pixel values (Schoenmakers et al., 2013). The encoding model approach has also been employed outside the visual domain, for example to characterize semantic representations of words (Huth et al., 2016). In this study, the feature space was defined as a basic dictionary of English words, and the model was fitted on whole-brain data. The high-dimensional representation of stimuli used in the encoding model approach typically translates into design matrices encompassing more predictors than time points. In this setting, a unique model fit can only be obtained by adding a regularization term to the model.

Model regularization adds additional constraints to the model fitting procedure on top of minimizing the error term. As the number of predictors approaches the number of collected time points (i.e. the length of the fMRI time series), model regularization becomes increasingly relevant, and for saturated models, regularization is indispensable. In the context of fMRI activation analyses, model regularization can improve the robustness of single-trial and FIR models, and is crucial for the encoding model approach.

For linear models, the two most common types of regularization are L1-regularization and L2-regularization (also known as lasso and ridge regression, Tibshirani, 1996; Hoerl and Kennard, 1970). While L1-regularization puts a threshold on the sum of absolute values of the beta estimates, L2-regularization bounds the sum of squared beta-values. These two types of regularization can result in fundamentally different beta estimates: L2-regularized models return nonzero beta-values for all predictors, whereas L1-regularized models return a sparse model fit, that is, most of the beta-values are set to zero and only a few predictors are included in the model fit. Whether to employ L1- or L2-regularization depends on a-priori assumptions on the data at hand. L2-regularization assumes that most of the predictors have an impact on the fMRI signal. In contrast, L1-regularization is based on the assumption that the fMRI signal can be modeled by a small fraction of the included predictors.

While both types of regularization have been employed in fMRI studies (Huth et al., 2016; Nishimoto et al., 2011), L2-regularization seems to occur more frequently in the neuroimaging literature than L1-regularization. This might be partly explained by the fact that fitting L1-regularized models is considerably more expensive, in terms of computation time, than fitting L2-regularized or unregularized models. While L2-regularized and unregularized models can be estimated using closed-form solutions, L1-regularization requires an iterative fitting procedure. Thus, estimating L1-regularized models instead of L2- or unregularized models substantially increases the running time of fMRI analyses. For certain types of analyses, for example whole-brain analyses on large samples, it is virtually infeasible to employ L1-regularization.

Here, we aim to facilitate the estimation of L1-regularized models on fMRI data. In the following, we present a package of functions for estimating L1-regularized models that are optimized for the mass-univariate approach. We describe the implementation of the functions, how to set their parameters, and provide benchmark results for two exemplary data settings.

## Methods

In the following, we assume to have an fMRI data matrix *Y* of size *n* × *ν*, with *n* being the number of time points and *ν* being the number of time series (e.g. the number of voxels). Moreover, we have a design matrix *X* of size *n* × *p*, with *p* being the number of predictors. The beta-values are stored in a matrix *B* of size *p* × *ν*. The intercept of the model is of size *n* × 1 and is denoted *I*, and the intercept’s beta-values are stored in *B*^0^ of size 1 × *ν*. For a vector *Z* of size *n* × 1, we define the mean squared error as 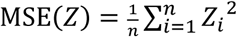. Matrix columns are indexed by *j*. We assume that the columns of the design matrix *X* are z-scored, i.e. 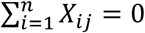 and MSE(*X*_*j*_) = 1 for all *j* = {1, …, *p*}. For a given regularization parameter *λ* ≥ 0, the L1-regularized model fit 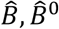 is expected to minimize, for all *j* = {1, …, *ν*}, the following objective function:

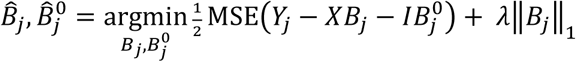

In contrast to L2-regularized or unregularized models, L1-regularized model fits cannot be computed using a closed-form solution. Instead, an iterative procedure is required to find 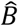. In the following sections, we describe two different algorithms for fitting L1-regularized models, coordinate descent (Friedman et al., 2010) and alternating direction method of multipliers (ADMM, Boyd et al., 2010), and how we optimized and implemented these algorithms for mass-univariate analyses.

### The CPU-based implementation: lasso_mex

In the CPU-based implementation, the model fit is computed using the coordinate descent algorithm proposed by Friedman et al., 2010. In each step of the iterative fitting procedure, the beta-value of a single predictor is updated while the remaining beta-values are fixed. As suggested by Friedman et al., 2010, the beta-values are computed using covariance updates, i.e. without explicitly computing residual values.

Friedman et al., 2010 suggest to compute covariances between predictors dynamically as required during the estimation process. However, in the mass-univariate setting, the same design matrix is used to model a large number of fMRI time series. Thus, instead of computing covariances between predictors dynamically for each voxel, we precompute the full covariance matrix of the design matrix before starting the coordinate descent, thereby avoiding redundant computations of predictor covariances across voxels.

Aside from precomputing the covariance matrix, our implementation includes features such as warm starts and active sets described in Friedman et al., 2010. The idea behind active sets is to iterate only through predictors whose beta-values were set to nonzero values at an earlier stage. Once convergence among the beta-values included in the active set is achieved, the algorithm iterates through all beta-values to check whether additional predictors have to be included. Using active sets can considerably speed up the estimation procedure, and moreover beta-values can be stored in sparse format, thereby reducing memory usage.

The estimation procedure for a given *λ* parameter can be considerably accelerated by properly initializing the beta-values, instead of starting with all beta-values set to zero. Such an initialization can be obtained from a prior estimate using a larger *λ* parameter. Generally, computation times increase as *λ* becomes smaller, due to the larger number of nonzero beta-values. Thus, successively fitting models along a decreasing sequence of lambda parameters using warm starts is typically faster than starting from scratch for each lambda value (Friedman et al., 2010).

To further accelerate the estimation procedure, the CPU-based implementation distributes computations among multiple CPU cores using a parallel for-loop over the voxels, exploiting the fact that models are fitted independently across voxels in the mass-univariate analysis approach. To this end, the algorithm was implemented in C++ using OpenMP for parallelization. The performance improvement achieved by this parallelization step depends on the number of available CPU cores.

### How to use lasso_mex

While the underlying coordinate descent algorithm is implemented in C++, the `lasso_mex` function can be conveniently called from Matlab via the mex API. The function takes a design matrix `X`, a matrix `Y` containing fMRI time series and a sequence of lambda-parameters `lambda_seq` as input. Technical parameters can be optionally specified using an `options` structure, otherwise default values are used. The columns of the design matrix `X` must be z-scored, and `X` must not contain an intercept column. The function returns beta-values in sparse format, with `b_values` containing the actual beta values, `b_indexes` containing the indexes of the values, and `N_nz` containing the number of nonzero beta-values for each voxel. Unregularized intercept coefficients are returned in `b0`.

~~~
[b_values, b_indexes, N_nz, b0] = lasso_mex(X, Y, lambda_seq);
~~~

The function `lasso_mex` is written in Matlab and calls, after some sanity checks and precomputations, the MEX function lasso_mex_cpp.c, which is written in C++ and runs the coordinate descent algorithm.

The resulting beta-values in sparse format can be converted to full format using the `convert_betas_sparse_to_full` function:

~~~
b_full = convert_betas_sparse_to_full(b_values, b_indexes, N_nz, size(X,2));
~~~

The sequence of lambda-parameters `lambda_seq` should be decreasing in order to benefit from warm starts as described above. Lambda-values are typically defined on a log-scale, e.g. *λ* ∈ {2^*k*^, 2^*k*−1^, …, 2^*k*−*n*^}. The lambda parameter determines the degree of model regularization. For lambda-values larger than a certain data-dependent threshold, all beta-values are set to zero, and the model fit only consists of the intercept. The critical threshold can be computed using the `calculate_lambda_start` function.

~~~
lambda_start = calculate_lambda_start(X, Y);
~~~

For lambda-values larger than `lambda_start`, all beta-values will be zero. Thus, the first value of the lambda sequence is typically set to `lambda_start` or to value from a predefined discrete scale that is close to `lambda_start`. How to set the smallest value of the lambda-sequence is less clear and involves a trade-off between computation time and the desired degree of model saturation. Generally, estimating weakly regularized models requires longer computation times than estimating strongly regularized models. As the number of nonzero beta-values approaches the number of time points of the fMRI time series, model estimation can become time-consuming and unstable in overparameterized settings (i.e. when *p* > *n*). Thus, the smallest value of the lambda sequence is typically chosen to achieve a certain degree of model saturation while keeping computation time within reasonable bounds.

Optionally, technical parameters of the `lasso_mex` function can be set via the `options` structure. The default values are:

~~~
options.n_iter_max = 1e5;
options.tol_value = 1e-3;
options.buffer_factor = 3;
options.cpu_load_factor = 1;
[b_values, b_indexes, N_nz, b0] = lasso_mex(X, Y, lambda_seq, options);
~~~

The `n_iter_max` parameter defines an upper bound for the number of iterations performed by the coordinate descent algorithm. This parameter can be set to a larger value if the default value results in the error message `Max. iter. reached, no convergence!`. However, reaching the maximum number of iterations can also indicate that model regularization is too weak and more stringent regularization is required.

The `tol_value` parameter determines the precision of the estimated beta-values. The coordinate descent algorithm stops when 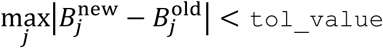. If higher than default precision is required, the `tol_value` parameter can be set to a smaller value, which will typically increase computation time. Vice versa, low-precision estimates can be obtained by setting `tol_value` to a larger value, which might accelerate the estimation procedure.

To minimize memory requirements, beta-values are stored in sparse format. The maximum number of nonzero beta-values per voxel is determined by the `buffer_factor` parameter. Internally, the `buffer_factor` is multiplied by *n* (the number of rows of the design matrix *X*) in order to compute how much memory is preallocated for the beta-values. In well-defined settings, i.e. if *p ≤ n*, the `buffer_factor` parameter can be set to 1. In overparameterized settings (i.e. if *p* > *n*), if the default value results in the error message `N_nz over maximum, larger buffer is required!`, the `buffer_factor` parameter should be set to larger value.

The parameter `cpu_load_factor` determines the degree of CPU utilization. For a `cpu_load_factor` of 1 (default value), all CPU cores are engaged, whereas for a `cpu_load_factor` of 0, only a single core is occupied. To distribute computations among all but one core, the `cpu_load_factor` can be set to 0.99.

### The GPU-based implementations: lasso_mexcuda and lasso_gpu

The two GPU-based implementations are based on the alternating direction method of multipliers (ADMM) algorithm described in Boyd et al., 2010. In contrast to the coordinate descent algorithm, the ADMM algorithm does not sequentially cycle trough the beta coefficients but instead all beta coefficients are updated simultaneously via matrix multiplication. While the ADMM procedure is less memory-efficient than coordinate descent, it can be accelerated on the GPU by parallelizing matrix multiplications and other steps. The first version (`lasso_mexcuda`) can be called from Matlab and does not require any Matlab toolboxes. After some sanity checks and precomputations, the mex functions ADMMcublasOverMex.c or ADMMcublasUnderMex.c are called (depending on whether the design matrix is overparameterized or not), which are written in C++. These functions then call ADMMcublasOver.cu or ADMMcublasUnder.cu, which contain CUDA code calling functions from the cuBLAS library to run the ADMM algorithm on the GPU. The second version (`lasso_gpu`) is implemented directly in Matlab using `gpuArray` and thus depends on the Parallel Computing Toolbox. For both versions, a CUDA-enabled GPU device is required (see also Table 1).

**Table 1.** Overview over hardware and software requirements of the different functions for estimating L1-regularized linear models. The `lasso` function is part of the Statistics and Machine Learning Toolbox and only included here for benchmarking purposes (see main text). The `lasso_mex`, `lasso_mexcuda` and `lasso_gpu` functions introduced here are specifically optimized for the mass-univariate analysis approach and depend on different hardware and software configurations.

| Function | Hardware requirements | Software requirements | Algorithm | Implementation |
| --- | --- | --- | --- | --- |
| `lasso` | CPU | Matlab, Statistics and ML Toolbox | Coordinate descent | Matlab |
| `lasso_mex` | CPU | Matlab | Coordinate descent | C++, OpenMP |
| `lasso_mexcuda` | CPU + CUDA-enabled GPU | Matlab, CUDA Toolkit | ADMM | C++, cuBLAS |
| `lasso_gpu` | CPU + CUDA-enabled GPU | Matlab, Parallel Computing Toolbox | ADMM | Matlab |

Both `lasso_mexcuda` and `lasso_gpu` use warm starts along the supplied lambda sequence, and the same considerations on the choice of the lambda sequence as discussed above in the `lasso_mex` section also apply to the `lasso_mexcuda` and `lasso_gpu` functions.

### How to use lasso_mexcuda and lasso_gpu

The functions take a design matrix `X`, a matrix `Y` containing fMRI time series and a sequence of lambda-parameters `lambda_seq` as input. Technical parameters can be optionally specified using an `options` structure, otherwise default values are used. The columns of the design matrix `X` must be z-scored, and `X` must not contain an intercept column. Beta-values are returned in full format in `B`, and unregularized intercept coefficients are returned in `B0`. As `lasso_mexcuda` and `lasso_gpu` are optimized for the GPU, `B` and `B0` are returned in single-precision format.

~~~
[B, B0] = lasso_mexcuda(X, Y, lambda_seq);
[B, B0] = lasso_gpu(X, Y, lambda_seq);
~~~

Optionally, the following technical parameters can be specified (set to default values here):

~~~
options.n_iter_max = 1e5;
options.tol_value = 1e-3;
options.buffer_size = 8192;
[B, B0] = lasso_mexcuda(X, Y, lambda_seq, options);
[B, B0] = lasso_gpu(X, Y, lambda_seq, options);
~~~

The `n_iter_max` parameter defines an upper bound for the number of iterations performed by the ADMM algorithm. This parameter can be set to a larger value if the default value results in the error message `Max. iter. reached, no convergence!`. However, reaching the maximum number of iterations can also indicate that model regularization is too weak and more stringent regularization is required.

The `tol_value` parameter determines the precision of the estimated beta-values. If higher than default precision is required, the `tol_value` parameter can be set to a smaller value, which will typically increase computation time. Vice versa, low-precision estimates can be obtained by setting `tol_value` to a larger value, which might accelerate the estimation procedure.

The `buffer_size` determines how many voxels are simultaneously processed on the GPU. Setting this parameter to a larger value might accelerate the estimation procedure, provided that the GPU has sufficient memory resources. If `buffer_size` exceeds the memory capacity of the GPU, Matlab terminates the function call with an error message.

### Benchmarking

The three functions `lasso_mex`, `lasso_mexcuda` and `lasso_gpu` were benchmarked in two data settings to demonstrate the efficiency of the implementations. In both benchmarks, *X* and *Y* data were randomly drawn from the normal distribution. Benchmark A corresponds to an overparameterized model, representing for example an encoding model with a large feature space. The number of time points was set to *n* = 300, which corresponds to a 10 min scanner run for a TR of 2 s. The number of predictors was set to *p* = 5000, i.e. *p* ≫ *n*. The model was estimated on *ν* = 65536 voxels, approximately corresponding to whole-brain data for an isotropic 3 mm resolution. The lambda sequence was set to {2^−2^, 2^−3^, …, 2^−6^}. Benchmark B corresponds to a well-defined setting, e.g. a single-trial or FIR model. The number of time points and voxels in setting B were identical to setting A, but the number of predictors was set to *p* = 200, i.e. *p* < *n*. The models were estimated for a single lambda-value *λ* = 2^−4^ to allow for a comparison of the computation times with L2-regularized and unregularized model fits.

The three functions `lasso_mex`, `lasso_mexcuda` and `lasso_gpu` were compared to the `lasso` function that is part of Matlab’s Statistics and Machine Learning Toolbox. The `lasso` function was repeatedly called to fit models for all voxels using a for-loop, taking a single time series as input in each call.

The benchmarks were run on a workstation equipped with a dual Intel Xeon E5-2665 CPU (16 cores overall), 32 GB memory and an Nvidia Quadro P2000 GPU. The functions were benchmarked on Windows 10 64-bit and Matlab R2019b, as well as Ubuntu 18.04 LTS and Matlab R2018b. On Windows, `lasso_mex` was compiled using mex and the MSVC compiler of Visual Studio 2017, and `lasso_mexcuda` using CUDA Toolkit 10.1.243. On Linux, `lasso_mex` was compiled using mex and gcc 6.5, and `lasso_mexcuda` using CUDA Toolkit 9.1.85.

Moreover, to assess the impact of the GPU hardware, we compared two different GPU devices using benchmark A, Nvidia’s Quadro P2000 (1024 cores, 5 GB memory) and Tesla V100 SXM2 (5120 cores, 16 GB memory). To this end, the functions `lasso_mexcuda` and `lasso_gpu` were run on a p3.2xlarge instance on Amazon Web Services (AWS) using a Matlab Amazon Machine Image (AMI).

The package including compiled binaries and source code is available at https://git.io/JvUpi.

## Results

On both benchmarks A and B, the `lasso_mex` function was considerably faster than the standard `lasso` function using a single CPU core, showing that the coordinate descent algorithm underlying `lasso_mex` is efficiently implemented (see Figure 1). Setting the `lasso_mex` function to distribute computations across multiple CPU cores led to further reductions of computation time. The parallel implementation of the ADMM algorithm on the GPU (`lasso_mexcuda`, `lasso_gpu`) provided further acceleration for benchmark A, with `lasso_gpu` being considerably faster than `lasso_mexcuda`. In absolute numbers, a reduction of computation time from approximately 9 h to 5 min could be achieved on Windows using the `lasso_gpu` function, see Supplementary Table 1. As shown in benchmark B, L2-regularized (ridge regression) or unregularized (ordinary least squares, OLS) model estimation remains faster than accelerated L1-regularization.

**Figure 1.**
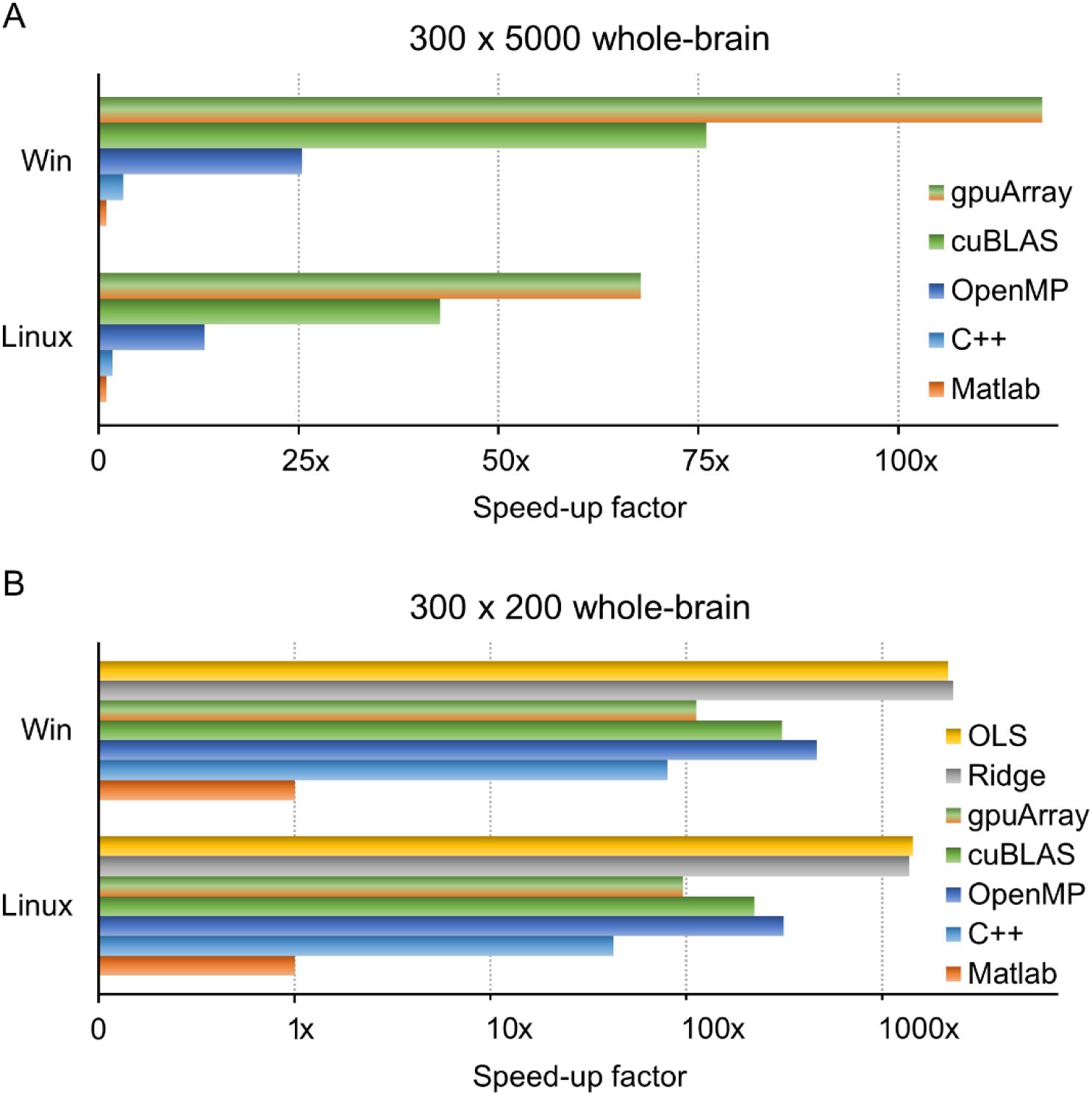
Benchmark results for the three lasso functions `lasso_mex`, `lasso_mexcuda` and `lasso_gpu`. **A:** Benchmark for an overparameterized setting with *n* = 300 time points and *p* = 5000 predictors, corresponding for example to an encoding model with a large feature space. The `lasso_mex` function required considerably less running time for whole-brain estimates than the standard `lasso` function (Matlab, orange) on a single CPU core (C++, light blue). Distributing computations across multiple CPU cores further reduced the running time of the `lasso_mex` function (OpenMP, dark blue). The `lasso_mexcuda` function, which runs the ADMM algorithm on a GPU using the cuBLAS library, further accelerated the estimation procedure (cuBLAS, green). The `lasso_gpu` function, which runs the ADMM algorithm on the GPU using the Parallel Computing Toolbox, provided the highest speed-up factor for benchmark A (gpuArray, green/orange). **B:** Benchmark for a well-defined setting with fewer predictors than time points (*n* = 300, *p* = 200), corresponding for example to a single-trial or FIR model. Again, on a single CPU core, the `lasso_mex` function fitted L1-regularized models more efficiently than the standard `lasso` function. Further speed-up could be achieved by distributing computations across multiple CPU cores. The GPU-based implementations (`lasso_mexcuda`, `lasso_gpu`) performed not as fast as the multicore CPU version on this benchmark, as the GPU was not fully occupied in this small-scale setting. Unregularized (OLS, yellow) or L2-regularized (ridge, gray) model estimation using closed-form solutions remains faster than accelerated L1-regularization. Absolute computation times are given in Supplementary Table 1.

The comparison of two different GPU devices using benchmark A on Linux revealed that the larger device (Tesla V100) achieves a 4.5x speed-up over the smaller card (Quadro P2000), see Figure 2. This acceleration approximately corresponds to the ratio of available CUDA cores of 5120:1024. In absolute numbers, computation time could be further reduced to 1 min using the `lasso_gpu` function on the V100 device, see Supplementary Table 2.

**Figure 2.**
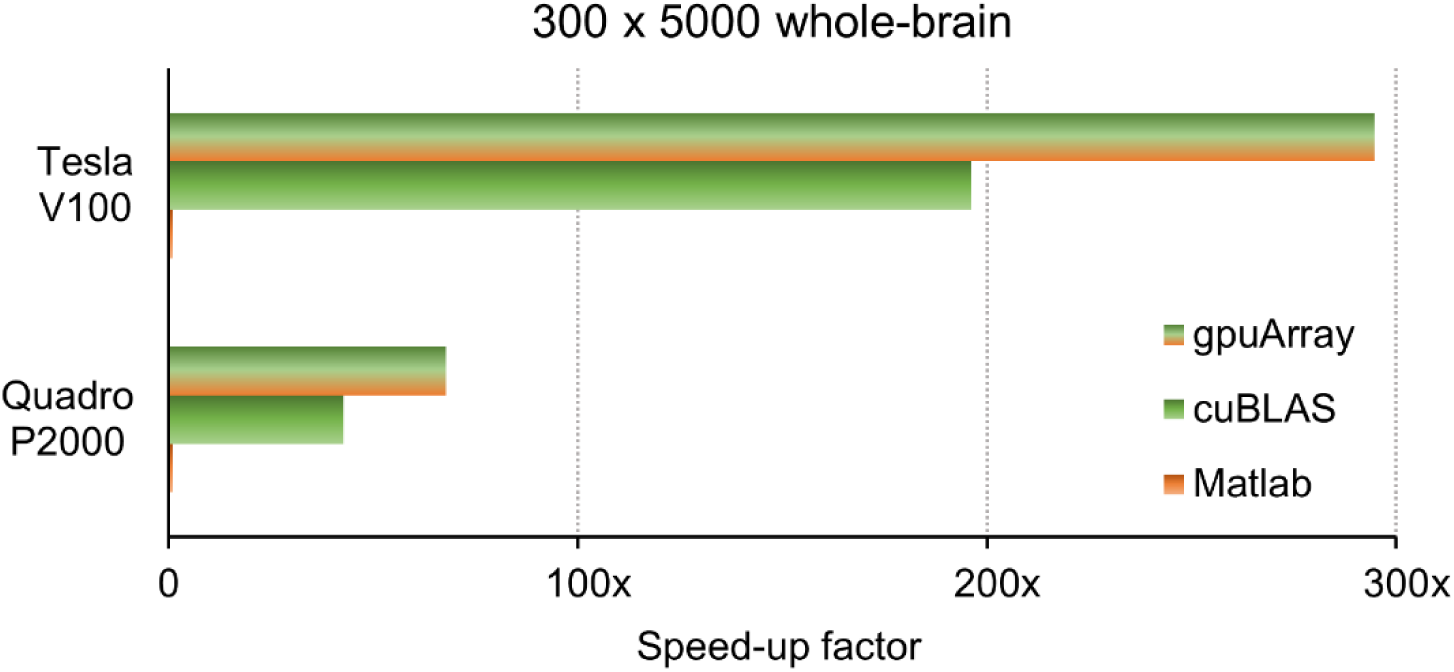
Benchmark results for two different GPU devices. Nvidia’s Tesla V100 GPU device yielded a speed-up of approximately 4.5x over the Quadro P2000 card, which roughly corresponds to the ratio of 5120:1024 cores. Absolute computation times are given in Supplementary Table 2.

## Discussion

We introduced a package of functions for estimating L1-regularized models in the mass-univariate fMRI analysis approach. The presented functions significantly accelerate the model estimation procedure and thereby facilitate the use of L1-regularization in the mass-univariate approach and enable its application on whole-brain data and large samples. While the presented benchmarks were performed on data corresponding to a single fMRI scanning session, efficiency gains scale up linearly for multiple sessions per subject and multiple subjects per sample. For example, for a sample of 20 subjects and 5 scanning sessions per subject, a computation time reduction from 9 hours to 5 minutes per scanning session translates into a reduction from 37 days to 8 hours for the whole sample.

This speed-up makes it practically feasible to employ L1-regularization in the context of the encoding model approach. While the majority of studies using the encoding model approach have been focused on visual processing (van Gerven, 2017), Huth et al., 2016 have shown that an extension to other domains such as whole-brain semantic representations is possible and promising. Data-driven generation of predictive features, for example via deep neural networks (Güclü and van Gerven, 2015; Kell et al., 2018; Mohr et al., 2019), typically results in large feature spaces and therefore requires efficient model estimation procedures. Given the rapid advancement of machine learning in a range of domains that are also relevant for neuroimaging (e.g. geometric representations of objects (Eslami et al., 2018) or spatial navigation (Banino et al., 2018)), we expect a proliferation of the encoding model approach for predicting fMRI signals via machine-learning generated features. Estimating such large-scale encoding models using L1-regularization can be efficiently performed by the functions presented here.

Moreover, the presented functions facilitate the use of L1-regularization for modeling approaches such as single-trial and FIR models (Mumford et al., 2012; Ollinger et al., 2001). To our knowledge, the impact of L1-regularization has not yet been systematically evaluated for these types of models. Using the functions presented here, future studies can efficiently evaluate how L1-regularization impacts the robustness and predictiveness of single-trial and FIR models in comparison to L2-regularization or unregularized model estimation.

Overall, the functions introduced here significantly accelerate L1-regularization in the mass-univariate setting and make it practically feasible to estimate L1-regularized models on whole-brain data and large samples.

## Acknowledgments

This work was supported by the German Research Foundation (DFG), grant CRC 940, project Z2. We thank the Center for Information Services and High Performance Computing (ZIH) at TU Dresden for generously providing computing resources.

## Supplementary Material

**Supplementary Table 1.** Computation times for benchmarks A and B in seconds.

| **300 x 5000** | **Windows** | **Linux** | **300 x 200** | **Windows** | **Linux** |
| --- | --- | --- | --- | --- | --- |
| OLS | - | - | OLS | 0.18 | 0.15 |
| Ridge | - | - | Ridge | 0.17 | 0.16 |
| gpuArray | 277.8 | 265.0 | gpuArray | 3.51 | 2.26 |
| cuBLAS | 431.6 | 420.5 | cuBLAS | 1.29 | 0.98 |
| OpenMP | 1290.8 | 1351.9 | OpenMP | 0.85 | 0.69 |
| C++ | 10567.6 | 10175.3 | C++ | 4.91 | 5.10 |
| Matlab | 32777.4 | 17952.3 | Matlab | 395.42 | 217.65 |

**Supplementary Table 2.** Computation times in seconds for benchmarks A on two different GPU devices.

| **300 x 5000** | **Tesla V100** | **Quadro P2000** |
| --- | --- | --- |
| gpuArray | 60.9 | 265.0 |
| cuBLAS | 91.5 | 420.5 |
| Matlab | 17952.3 | 17952.3 |

## Notes

https://git.io/JvUpi

